# Cooperation not competition: bihemispheric tDCS and fMRI show role for ipsilateral hemisphere in motor learning

**DOI:** 10.1101/082669

**Authors:** Sheena Waters, Tobias Wiestler, Jörn Diedrichsen

**Affiliations:** Institute of Cognitive Neuroscience, University College London, London, UK; Institute of Neurology, University College London, London, UK; Sobell Department for Motor Neuroscience and Motor Disorders, University College London,London UK; Brain and Mind Institute, Western University, London ON, Canada

## Abstract

What is the role of ipsilateral motor and pre-motor areas in motor learning? One view supposes that ipsilateral activity suppresses contralateral motor cortex, and thus needs to be inhibited to improve motor learning. Alternatively, the ipsilateral motor cortex may play an active role in the control and learning of unilateral hand movements. We approached this question by applying double-blind bihemispheric transcranial direct current stimulation (tDCS) over both contralateral and ipsilateral motor cortex in a between-group design during four days of unimanual explicit sequence training. Independently of whether the anode was placed over contralateral or ipsilateral motor cortex, bihemispheric stimulation yielded substantial performance gains relative to unihemispheric or sham stimulation. This performance advantage appeared to be supported by plastic changes in both hemispheres: First, we found that behavioral advantages generalized strongly to the untrained hand, suggesting that tDCS strengthened effector-independent representations. Secondly, functional imaging during speed-matched execution of trained sequences conducted 48 h after training revealed sustained, polarity-independent increases in activity in both motor cortices relative to the sham group. These results suggest a cooperative rather than competitive interaction of the two motor cortices during skill learning and suggest that bihemispheric brain stimulation during unimanual skill learning may be beneficial, because it harnesses plasticity in the ipsilateral hemisphere.

**Significance statement:** Many neurorehabilitation approaches are based on the idea that is beneficial to boost excitability in the contralateral hemisphere while attenuating that of the ipsilateral cortex to reduce interhemispheric inhibition. We observed that bihemispheric tDCS with the excitatory anode either over contralateral or ipsilateral motor cortex facilitated motor learning nearly twice as strongly as unihemispheric tDCS. These increases in motor learning were accompanied by increases in fMRI activation in both motor cortices that outlasted the stimulation period, as well as increased generalization to the untrained hand. Collectively, our findings suggest a cooperative—rather than competitive—role of the hemispheres and imply that it is most beneficial to harness plasticity in both hemispheres in neurorehabilitation of motor deficits.

## Introduction

Even strictly unilateral motor behaviors—such as moving one hand—rely on interaction between cerebral hemispheres. However, the nature of this interaction has been disputed (1). One influential idea is the interhemispheric competition model, according to which the main interaction between the hemispheres can be characterized by mutual suppression via inhibitory projections (2, 3). This idea has shaped the thinking about how to improve motor learning through brain stimulation: it has been suggested that hand function can be improved not only through excitatory stimulation of contralateral motor cortex, but also through inhibition of ipsilateral cortex (4-7). tDCS is a non-invasive brain stimulation method that modulates cortical excitability. Classically, tDCS has been shown to increase motor-evoked potentials (MEPs) elicited by transcranial magnetic stimulation when an anode is placed above primary motor cortex (M1) (8) and to decrease MEPs in the presence of a cathode over M1 (9). Consistent with the interhemispheric competition model, Vines, Cerruti and Schlaug (10) demonstrated that bihemispheric tDCS—with the cathode located over ipsilateral M1—accelerates skilled motor performance more than unihemispheric tDCS. The idea of promoting motor learning by reducing ipsilateral excitability is particularly pertinent to stroke neurorehabilitation, where it has led to the supposition that suppressing activity in the healthy hemisphere to release inhibition of lesioned cortex may facilitate recovery (6, 11-17).

There are several reasons to doubt the competition model as an explanation of the bihemispheric stimulation advantage. First, it is unclear whether cortical excitability is consistently suppressed below the cathode. For example, during 2 mA tDCS stimulation, MEP magnitude underneath the cathode appears to increase rather than decrease (18). Secondly, while consistent MEP decreases have been measured under the cathode after unihemispheric stimulation (9), smaller or no polarity-specific excitability changes have been found after bihemispheric stimulation (19). Collectively, these findings raise the possibility that increased neural plasticity after tDCS stimulation is not related to the polarity-specific excitability changes that can be measured with TMS, but rather due to the electrical current running transversely through the cortical tissue (20). If this were true, then tDCS effects on plasticity should only depend on the spatial distribution of these currents, but not on their direction.

Rather than simply inhibiting the contralateral cortex, the ipsilateral motor cortex may have an active role in the execution and learning of complex movements. While unilateral movement deactivates ipsilateral M1 relative to rest (21, 22), recent functional magnetic resonance imaging (fMRI) (22, 23), and electrophysiological (24) studies have demonstrated that activity in M1 contains detailed information about ongoing ipsilateral movements. This activity could contribute to movement control either through directly descending ipsilateral projections, or by shaping activation patterns on the contralateral side (25). According to this ‘interhemispheric cooperation’ model, bihemispheric tDCS is more effective than unihemispheric stimulation not because it silences ipsilateral cortex, but rather because it actually increases ipsilateral plasticity.

To adjudicate between these two explanations for the superior efficacy of bihemispheric tDCS, we trained 64 participants in a double-blind study, either using sham, unihemispheric, conventional bihemispheric, or reversed-polarity bihemispheric tDCS (**Fig.1a**). Participants were trained on an unimanual sequence task on either the left or right hand over four days (**Fig. 1b**). After training, participants underwent a behavioral post-test and fMRI without tDCS to elucidate the neural changes induced by the stimulation.

**Figure 1:**
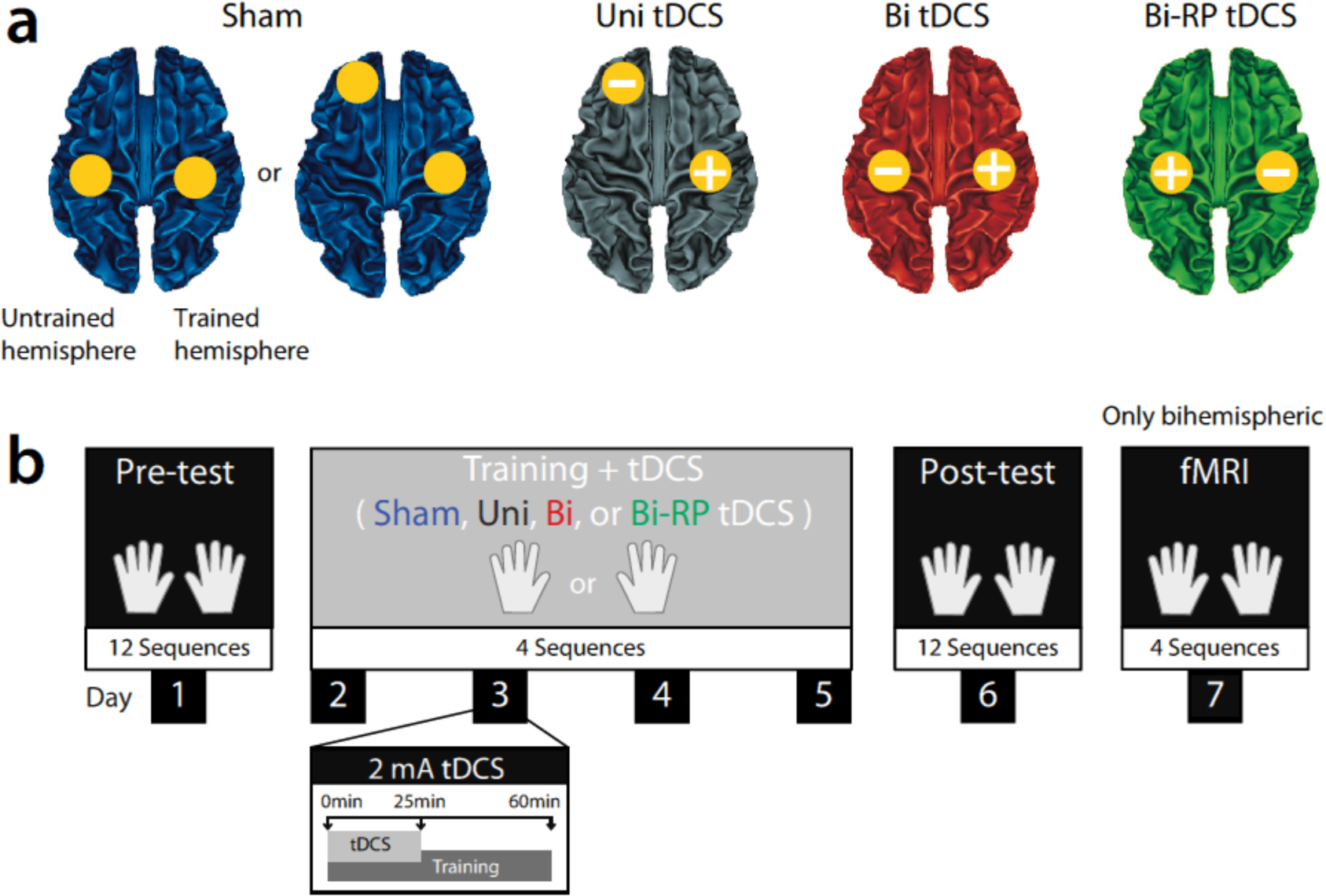
Experimental Design. **(a)** tDCS montages: sham stimulation (blue); unihemispheric (‘Uni’) tDCS (grey); bihemispheric (‘Bi’) tDCS (red); and reversed-polarity bihemispheric (‘Bi-RP’) tDCS (green). The location of the anode is indicated by a ‘+’ and the location of the cathode by a ‘−’. We show here the electrode placement for the left-hand trained group, for which the right hemisphere was the “trained hemisphere”. For right-hand trained participants, the electrode location was reversed for all stimulation groups. For purposes of double-blinding, 2/3 of the sham group had a bihemispheric, 1/3 unihemispheric montage. (b) Procedure: During pre-test, participants performed 12 5-digit sequences with either hand. Subsequently, participants were assigned to one of the four tDCS groups and trained for 4 days with either the left or the right hand, resulting in 8 different groups (for details see **Table S1**). During the post-test, participants were tested again as in the pre-test without tDCS. Finally, all participants with a bihemispheric montage (but not with unihemispheric montage) underwent functional MRI scanning on day 7.

The interhemispheric competition model predicts that the reversed-polarity bihemispheric montage should impair performance even beyond that of the sham group (4, 6, 7, 10). Not only should contralateral cathodal stimulation suppress the motor areas involved in learning, but ipsilateral anodal stimulation should further increase interhemispheric inhibition. Additionally, functional imaging should reveal opposite changes in the hemisphere that received anodal and cathodal stimulation.

In contrast, the interhemispheric cooperation model predicts that plasticity is induced regardless of current direction, and that the behavioral advantage arises from plasticity in both hemispheres. Therefore, reversed-polarity tDCS should be as effective as conventional bihemispheric tDCS, and both should be more effective than the unihemispheric montage. Given the bilateral nature of the predicted plastic changes, the tDCS induced advantages should generalize to the untrained hand. Finally, bihemispheric tDCS should lead to the activity changes in both hemispheres in a polarity-unspecific manner.

## Results

### Bihemispheric tDCS increases motor learning more than unihemispheric tDCS

We first determined whether conventional bihemispheric tDCS is more effective in promoting learning than unihemispheric stimulation. We used the error-adjusted execution time (ET, see methods) as an overall performance measure. By the end of 4 days of sequence training, unihemispheric tDCS recipients executed sequences 0.203 ± 0.101 s (16.6%) faster than sham (*F*_(1,31)_ = 4.43, *p* = 0.043; **Fig.2a**, *black vs. blue lines).* As in previous reports (26), this advantage persisted during the post-test, which was conducted without tDCS one day after the final training session (*F*_(1,31)_ = 8.13, *p* = 0.008). There was no significant difference between the left and right hand trained groups in the size of this effect (*F*_(1,31)_ = 2.03, *p* = 0.164).

**Figure 2:**
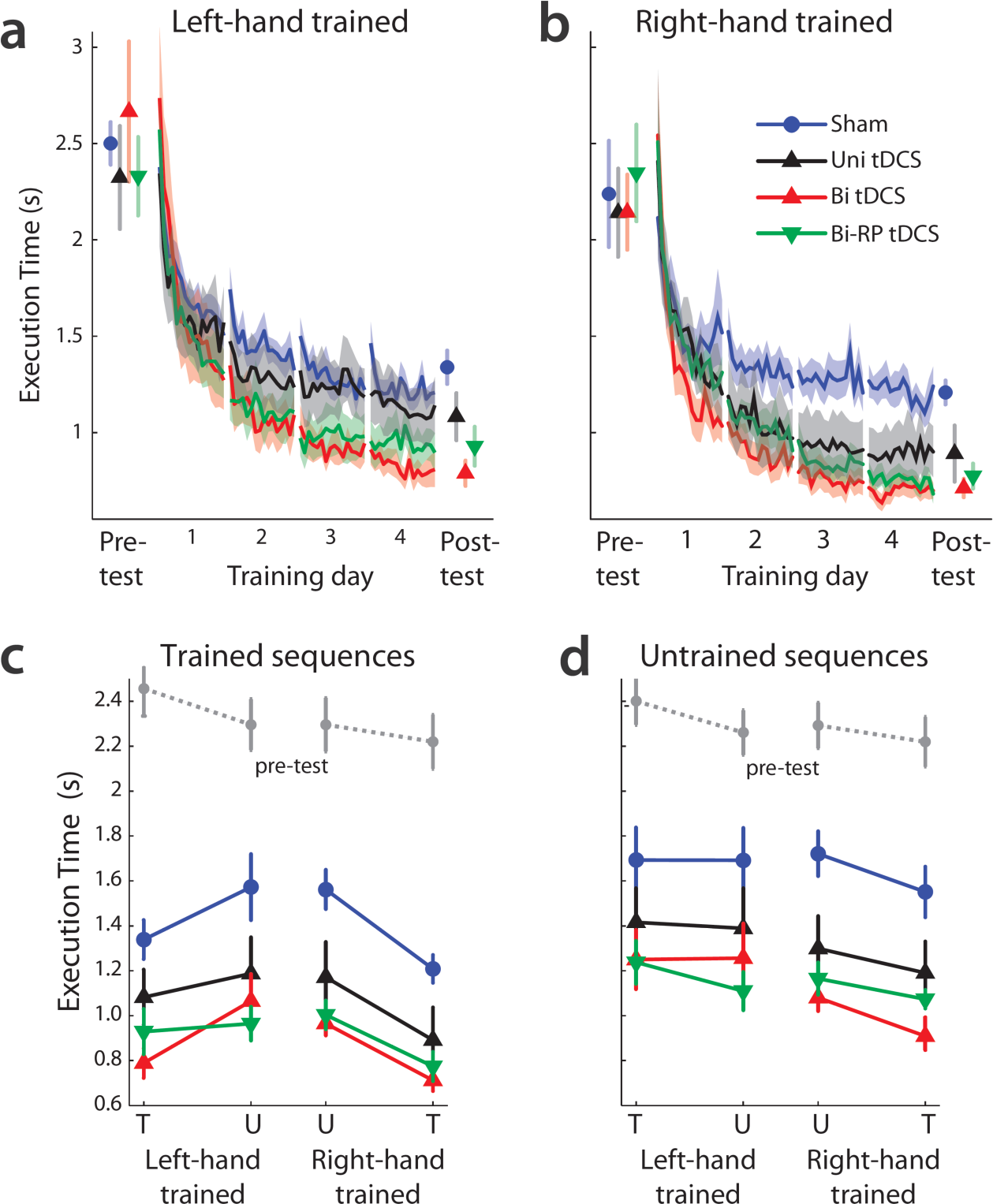
Bihemispheric tDCS accelerates learning and generalization in a polarity-independent manner. (**a**) Average ET in pre-test, training, and post-test for sham (blue), unihemispheric (black), bihemispheric (red), and reversed-polarity bihemispheric (green) tDCS groups. Subjects trained with either the left hand or (**b**) right hand. (**c**) Pre- and post-test data for the trained and (**d**) untrained sequences. Results are shown for the trained (T) and untrained (U) hands, separated by group. The pre-test results (grey dashed line) are averaged across all four groups. Error bars and shaded region indicate between-subject SEM.

Participants that received conventional bihemispheric tDCS were an *additional* 0.260 ± 0.114s faster than those who received unihemispheric stimulation (*F*_(1,24)_ = 12.44, *p* = 0.002; **Fig.2a**, bihemispheric vs. sham: *F*_(1,30)_ = 64.02, *p* = 6.24^−09^), which was maintained during the post-test (respectively, *F*_(1,24)_ = 10.98, *p* = 0.003 and *F*_(1,30)_ = 55.302, *p* = 2.75e^−08^). Hence, moving the cathode from the supraorbital ridge (unihemispheric tDCS) to ipsilateral M1 (bihemispheric tDCS) yielded nearly twice the performance gain (37.8% relative to sham). Therefore, we replicate here higher effectiveness for bihemispheric tDCS relative to unihemispheric stimulation (10, 27-31) in the context of a multiple-day learning study.

### Reversed-polarity bihemispheric tDCS increases motor learning

The interhemispheric competition model supposes that the advantage of bihemispheric relative to unihemispheric tDCS arises due to cathodal suppression of ipsilateral cortex. This idea predicts that the reversed-polarity montage should attenuate motor learning relative to sham, as excitability of contralateral M1 decreases both directly through cathodal and indirectly due to ipsilateral anodal stimulation.

Our data, however, showed the converse: reversed-polarity bihemispheric tDCS not only led to significantly greater ET advantages than sham, but was on par with conventional bihemispheric tDCS (**Fig.2a,b**). During the final training day, reversed-polarity tDCS elicited 0.376 ± 0.072 s (30.7%) faster ETs than sham (*F*_(1,31)_ = 27.55, *p* = 1.154e^−05^), but was not different from conventional bihemispheric tDCS (*F*_(1,23)_ = 1.81, *p* = 0.191). Similar findings held true for post-test performance: reversed-polarity bihemispheric tDCS led to faster ETs than sham (*F*_(1,30)_ = 25.0, *p* = 2.32e^−05^) and similar ETs to conventional bihemispheric tDCS (*F*_(1,23)_ = 1.93, *p* = 0.178). The trained hand advantages for the conventional and reversed-polarity bihemispheric groups were symmetric across left and right hand trained groups: the tDCS group x hand cohort interaction was not significant (*F*_(1,24)_ =0.11, *p* = 0.7459).

Collectively, these results indicate that bihemispheric stimulation is more effective than unihemispheric stimulation. This additional benefit, however, is not conferred by ipsilateral suppression, as we did not find a polarity-specific effect for bihemispheric tDCS. Instead, *any* stimulation of the ipsilateral motor areas accelerated motor learning.

These results, therefore, favour the bihemispheric cooperation model, by which the additional learning advantage of bihemispheric relative to unihemispheric tDCS arises from plastic changes in both hemispheres that would promote performance for both hands. This idea makes two testable predictions. First, a considerable part of the behavioral tDCS advantage should generalize to the untrained hand. Secondly, neural changes should occur in both hemispheres in a polarity-unspecific manner. In the remainder of the paper, we test these predictions.

### Behavioral tDCS effects generalize to the untrained hand

In the pre-test and post-test, participants performed trained sequences with their untrained hand, allowing us to assess inter-manual generalization. Consistent with previous results (32, 33), even the sham group showed considerable performance improvements on the untrained hand (**Fig.2c**). This generalization was even more pronounced in the tDCS groups: relative to sham, unihemispheric (*F*_(1,31)_ = 12.43, *p* = 0.001), bihemispheric (*F*_(1,24)_ = 26.99, p = 1.34e^−05^), and reversed-polarity bihemispheric groups (*F*_(1,24)_ = 25.98, p = 1.78e^−05^) all performed significantly better on the untrained hand. Additionally, performance for the untrained hand was better in bihemispheric relative to unihemispheric tDCS recipients (*F*_(1,31)_ = 4.26, *p* = 0.05), and there was no difference between the bihemispheric tDCS groups (*F*_(1,23)_ = 0.05, *p* = 0.822).

Importantly, the enhancement of untrained hand performance was even larger than what would have been expected if the generalization was just proportional to the improvements on the trained hand: when expressing the pre-/post-test difference for the untrained hand relative the learning gains for the trained hand, we found that tDCS increased the proportion of generalization. For sham recipients, the untrained hand gained 58.5% of the improvement of the trained hand, whereas, for both bihemispheric tDCS groups, the percentage of inter-manual generalization was greater (86-89.2%; *t*_(1,33)_ > 2.662, *p* < 0.012). These results suggest that bihemispheric tDCS influenced mainly effector-independent representations in both hemispheres.

### Bihemispheric tDCS causes activation increases in both hemispheres

To elucidate the neural consequences of tDCS stimulation, we measured fMRI BOLD activation while participants executed the four sequences with either the trained or untrained hand. Participants were scanned two days after their final tDCS-coupled training session, such that our measure would reflect learning-related plasticity, rather than immediate effects or aftereffects of tDCS on neural excitability or hemodynamics which might not be directly related to sequence learning (34-36). To ascertain that activity differences were not due simple behavioural differences, we attempted to closely match the performance in the scanner in terms of movement speed and force across both groups and hands. The hemispheric cooperation model predicts that similar neural changes should occur in both hemispheres, independently of polarity. As expected, for trained hand executions, the sham group exhibited contralateral activation and ipsilateral deactivation in M1 and S1 (**Fig. 3a**, **Fig. S1**), as often observed during simple unimanual hand movements such as those that we study here (22, 23). In contrast, both bihemispheric tDCS groups—independently of polarity—showed greater contralateral activation and no ipsilateral deactivation.

**Figure 3:**
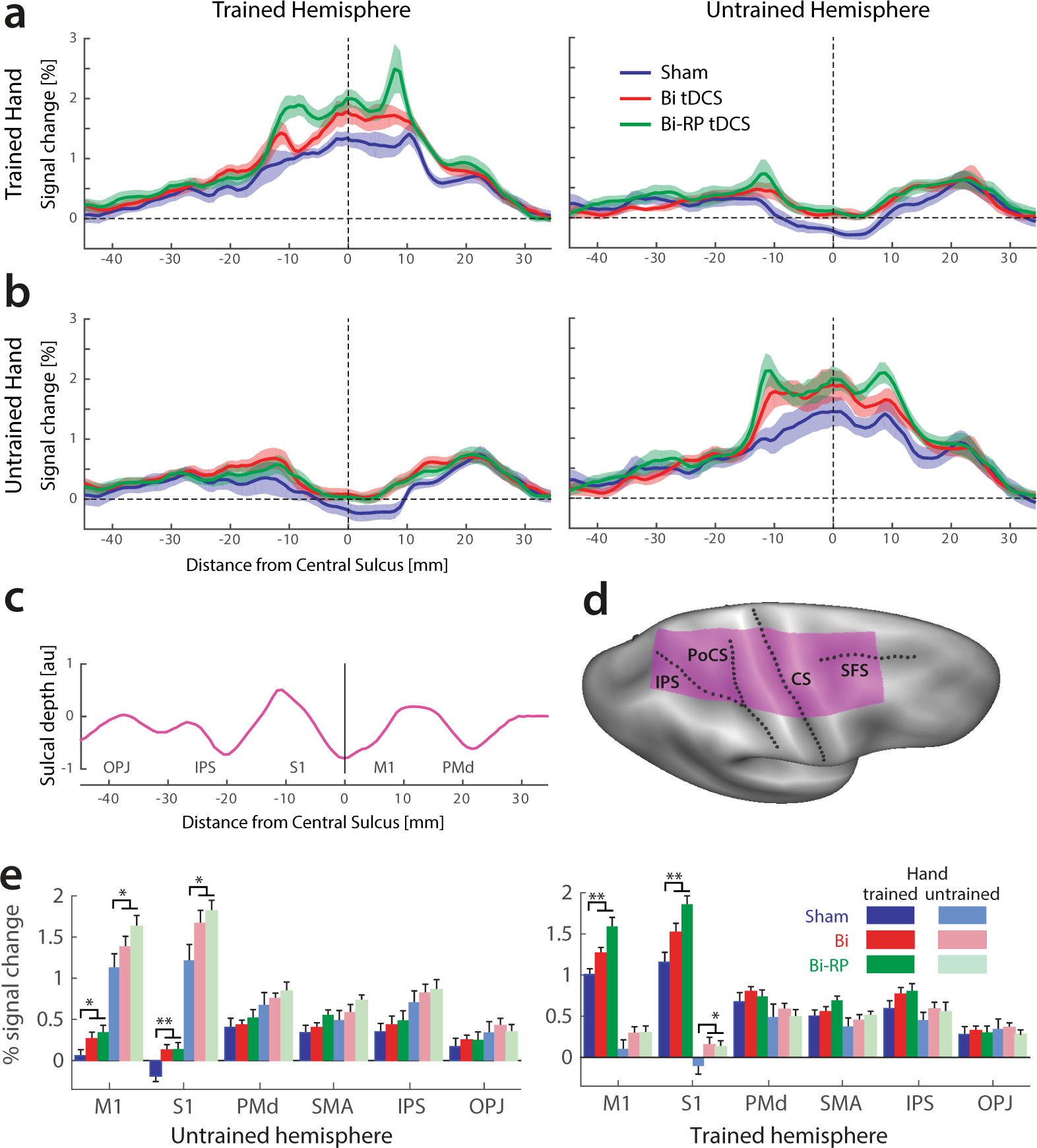
tDCS recipients show greater average activation than the sham group in sensorimotor areas of both hemispheres. **(a)** Profile plot of % signal change relative to rest in the trained and untrained hemispheres for trained hand executions. Results are shown for the sham, bihemispheric (Bi tDCS) and reversed-polarity tDCS groups (Bi-RP tDCS). The x-coordinate indicates the distance from the fundus of the central sulcus along the cortical surface in mm, running from occipital-parietal junction (posterior) to the rostral tip of premotor cortex (anterior). **(b)** Profile plot for the untrained hand. **(c)** Average sulcal depth along the profile. **(d)** Location of areas averaged in the profile plots on an inflated brain surface (purple area) **(e)** Average % signal change in 6 anatomically-defined ROIs (22, 32). Brackets indicate that the two bihemispheric groups were significantly different form sham (**: *p*<0.0083, statistical threshold for multiple comparisons, *: *p*<0.05). Abbreviations: CS = central sulcus; IPS = intraparietal sulcus; PoSC = postcentral sulcus; SFS = superior frontal sulcus.

For statistical comparison, we combined the two bihemispheric stimulation groups into a single tDCS group, as we did not find any significant clusters of differential activation between conventional and reversed-polarity tDCS groups. A surface-based group analysis (see Materials and methods) showed that both contralateral and ipsilateral primary somatosensory (S1) and primary motor (M1) cortex had greater activation in tDCS recipients (**Table 1**) for movements of the trained hand. Functional differences for the untrained hand were visually similar, but statistically less pronounced (**Fig.3b**; **Fig.S1b**; **Table 1**).

**Table 1:** Effect of tDCS on average activation using surface-based analysis. Between-subject analysis with uncorrected height threshold of *T*_(1,39)_ = 2.71, *p* = 0.005. Area indicates the size of the suprathreshold cluster, *T*_(1,39)_ the maximal *t*-value, *P*-cluster is the corrected probability of observing a cluster of this size or bigger over the whole cortical surface by random chance (52). Coordinates reflect the location of the cluster peak in MNI space.

| Region (Brodmann Area) | Area (mm<sup>2</sup>) | Peak value T<sub>(1,39)</sub> | P (clust) | MNI coordinates |  |  |
| --- | --- | --- | --- | --- | --- | --- |
|  |  |  |  | x | y | z |
| **<u>AVERAGE ACTIVATION</u>** |  |  |  |  |  |  |
| **tDCS>sham (% signal activation > rest)** |  |  |  |  |  |  |
| **<u>Trained hand</u>** |  |  |  |  |  |  |
| **Contralateral (Trained) hemisphere** |  |  |  |  |  |  |
| Postcentral (S1; BA3) | 145.33 | 3.64 | 0.007 | 49.89 | -22.22 | 51.78 |
| Postcentral (S1; BA3) | 116.21 | 3.81 | 0.030 | 18.26 | -39.58 | 65.61 |
| Precentral (M1; BA4) | 114.01 | 3.78 | 0.033 | 37.77 | -20.16 | 57.22 |
| **Ipsilateral (Untrained) hemisphere** |  |  |  |  |  |  |
| Precentral (M1; BA4) | 574.01 | 5.11 | <0.001 | -37.41 | -26.36 | 55.81 |
| Postcentral (S1; BA3) | 229.65 | 4.45 | <0.001 | -17.03 | -39.31 | 69.61 |
| Orbital area / Pars triangularis (BA47) | 182.63 | 5.49 | 0.002 | -42.46 | 29.98 | -0.78 |
| **<u>Untrained hand</u>** |  |  |  |  |  |  |
| **Contralateral (Untrained) hemisphere** |  |  |  |  |  |  |
| Orbital area / Pars triangularis (BA47) | 116.31 | 4.39 | 0.031 | -41.68 | 30.01 | -0.44 |
| **Ipsilateral (Trained) hemisphere** |  |  |  |  |  |  |
| *All non-significant* |  |  |  |  |  |  |

Analysis using anatomically predefined regions of interests (ROIs) (22), confirmed these results (**Fig. 3e**). Across both hemispheres and hands, we observed activation increases in M1 *(F*_(1,40)_ = 9.34, *p* = 0.004) and S1 (*F*_(1,40)_ = 13.55, *p* = 0.0007). For both regions, there was evidence for an interaction of tDCS x hemisphere x hand (M1: *F*_(1,40)_ = 3.72, *p* = 0.06; S1: *F*_(1,40)_ = 4.16, *p* = 0.048), reflecting that bihemispheric tDCS especially increased activation associated with the trained hand and ipsilateral (untrained) hemisphere.

Despite our best efforts to match behavioral performance during fMRI, there were slight but significant differences between the groups (**Table 2**). However, the differences in ET were very small (on the order of 40-60 ms), and the differences in force relative to sham were not consistent across the two bihemispheric tDCS groups (force was slightly lower than sham in the conventional bihemispheric tDCS group and slightly higher than sham in the reversed-polarity tDCS group). Nevertheless, to ensure that the observed increases in activation could not be caused by these small behavioral differences, we included execution time, error rate, and force as covariates in ANCOVA analyses and found that, for all comparisons, the effect of tDCS remained significant for both M1 (respectively, *F*_(1,39)_ = 7.26, *p* = 0.01; *F*_(1,39)_ = 14.03, *p* = 0.001; *F*_(1,39)_= 8.29, *p* = 0.006) and S1 (*F*_(1,39)_ = 8.02, *p* = 0.007; *F*_(1,39)_= 10.99, *p* = 0.002; *F*_(1,39)_= 12.78, *p* = 0.001).

**Table 2:** Execution time, error rate, and force during fMRI. Table shows mean (±standard error) of behavioral parameters for all three tDCS groups during fMRI scanning (averaged across hand training cohort). The last column indicates an *F*-test comparing the 3 groups.

|  | Sham |  | Bi tDCS |  | RP-Bi tDCS |  | ANOVA across groups |  |
| --- | --- | --- | --- | --- | --- | --- | --- | --- |
| | mean | SE | mean | SE | mean | SE | $F_{(2,39)}$ | $p$ |
| **<u>Trained Hand</u>** |  |  |  |  |  |  |  |  |
| Execution time (s) | 1.36 | (0.03) | 1.30 | (0.02) | 1.32 | (0.02) | 2.602 | 0.0869 |
| Error rate (%) | 13.76 | (2.36) | 5.08 | (1.18) | 5.61 | (1.70) | 7.206 | 0.0022 |
| Force (N) | 5.99 | (0.34) | 5.65 | (0.29) | 6.85 | (0.22) | 4.627 | 0.0157 |
| **<u>Untrained Hand</u>** |  |  |  |  |  |  |  |  |
| Execution time (s) | 1.40 | (0.04) | 1.32 | (0.02) | 1.31 | (0.01) | 4.506 | 0.0174 |
| Error rate (%) | 15.43 | (2.03) | 6.50 | (1.23) | 5.75 | (1.30) | 11.864 | 0.0001 |
| Force (N) | 5.69 | (0.26) | 5.50 | (0.27) | 6.71 | (0.29) | 5.718 | 0.0066 |

Taken together, these data demonstrate that bihemispheric tDCS—irrespective of the polarity—was associated with increases in average activation in both ipsilateral and contralateral hemispheres.

## Discussion

Our study provides evidence for an active role of ipsilateral motor regions in unimanual motor skill learning. We replicated the observation that bihemispheric tDCS is more effective than stimulating only one hemisphere (10). Critically, bihemispheric tDCS with the anode over ipsilateral motor cortex led to similar learning advantages as with the anode over contralateral motor cortex. Finally, both montages led to long-lasting increases of functional activity in bilateral sensorimotor areas and the tDCS-induced learning advantage generalized to the untrained hand.

While other studies have demonstrated an active role of the ipsilateral cortex in unimanual motor control in older adults (37, 38), we provide here evidence in young, healthy individuals. Our results also argue clearly against the interhemispheric-competition model as an explanation for the advantage of bihemispheric over unihemispheric tDCS. According to this idea, cathodal stimulation suppresses the ipsilateral hemisphere, subsequently releasing the contralateral hemisphere from interhemispheric suppression. Our results for the reversed-polarity bihemispheric group is at odds with this explanation, because excitatory stimulation of the ipsilateral cortex should have led to attenuated, rather than accelerated, motor learning relative to unihemispheric and even sham stimulation.

Instead of polarity-specific excitability changes in cortex underlying the electrodes, we suggest that these tDCS effects can be best explained by the spatial distribution of electrical current. Biophysical current modelling of tDCS (31, 39) demonstrates that the unihemispheric montage primarily sends current through the contralateral premotor and ipsilateral prefrontal regions, whereas bihemispheric stimulation targets motor and premotor regions bilaterally. Importantly, the weak radial currents that give rise to the polarity-specific effects on MEPs switch directions on opposite sides of the gyrus (20). Therefore, premotor areas in both hemispheres could experience either suppression or excitation depending on their folding geometry. Rather, the effects of tDCS on neuroplasticity are more likely mediated by the considerably stronger tangential currents, which, in contrast to the radial currents, do not have polarity-specific effects (20). Thus, changes in MEPs and changes in plasticity may ultimately rely on different mechanisms, which would explain why bihemispheric stimulation elicits larger polarity-independent behavioral effects relative to unihemispheric tDCS (10), but smaller or no polarity-specific MEP changes (19).

Increased plasticity in motor and pre-motor areas of both hemispheres would also explain our observation that the behavioral advantages due to tDCS generalized to the untrained hand. In a previous study, we used multi-voxel pattern analysis to identify commonalities in the neural encoding of specific motor sequences for the left and right hands (32) We found widespread, shared representations in both premotor and, surprisingly, primary motor areas of both hemispheres. Given that tDCS stimulation increased the amount of intermanual generalization, it appears likely that tDCS especially boosted these effector-independent sequence representations.

Consistent with this idea, we found that the average activity during trained hand movements was larger in the bihemispheric tDCS groups in both contra- and ipsilateral sensorimotor regions. The results for the untrained hand were similar, albeit less robust. Previous work has demonstrated bilateral activity increases in M1 during the application of bihemispheric tDCS (29). Similar online (40, 41) or short-lasting (42-46) changes underneath the anode have also been reported for unihemispheric tDCS. Importantly, we measured functional activity approximately 48 hours after the end of the final stimulation. Therefore, the activity increases reported here likely reflect longer-lasting changes caused by neural plasticity, rather than any immediate effects of tDCS on the hemodynamic response. For example, it is possible that decreases of local inhibition within both motor cortices (47) during stimulation led to increased neural recruitment within primary motor cortex during the 4 days of sequence learning (48).

The fact that the activation changes in our study were restricted to primary motor and sensory cortex should not necessarily be taken as evidence that the plastic changes that underlie the behavioral effects also took place here. Rather, it is equally possible that tDCS led to increased plasticity bilaterally in premotor or supplementary motor cortex, and that the increased activity in M1 and S1 reflects the increased modulatory input from these areas.

To summarise, we demonstrate that conventional and reversed-polarity bihemispheric tDCS similarly increase motor learning and BOLD activation in both anode- and cathode-modulated hemispheres. tDCS effects are still often construed in terms of simple excitation/inhibition of neural tissue under the electrodes, and the interhemispheric competition model is commonly used to explain the effects of bihemispheric tDCS. Our results suggest that this model should be abandoned in favor of a framework that acknowledges the broad effects of tDCS current and the roles that both hemispheres play in the encoding of information during fine motor learning.

## Materials and methods

### Participants

Sixty-four healthy right-handed subjects (54.69% females; average age 22.84 ± 0.56 years) participated in this study. Participants completed the Edinburgh Handedness Inventory (49), as well as a survey of medical history, musical and computer gaming experience, and previous exposure to brain stimulation. At the beginning of each session, participants provided information about sleep quality, alertness, attention, and task difficulty using visual analogue scales ranging from 0 (lowest) to 10 (highest). There were no significant differences between stimulation groups in terms of these parameters (**Table 2**). The protocol was approved by the UCL Research Ethics Committee. Exclusion criteria for participation were as detailed previously (33). Participants gave written informed consent in accordance to the Declaration of Helsinki, and received financial remuneration with the possibility of additional bonuses for completing the study and performance improvements.

### Sequence task

The experiment required the fast production of different unimanual 5-finger sequences. The sequences were performed on a custom-built, MRI-compatible, piano-like isometric force keyboard. Forces exerted on each key were measured every 5 ms via transducers (Sensing & Control Honeywell Inc., FSG-15N1A, dynamic range of 0-25 N).

The sequence task required participants to press each finger in a pre-defined order. A computer screen showed each sequence as a specific ordering of the numbers 1-5. ‘1’ referred to left-most and ‘5’to right-most key, and the sequence was executed from left to right. In other words, sequences were cued in an extrinsic, spatial reference frame. Based on pilot experimentation (22), we selected twelve sequences of matched difficulty, excluding sequences that contained a run of more than three adjacent digits. From these twelve sequences, each participant was assigned a set of four sequences that would be trained (32).

A small (0.53cm × 0.53cm) green box flanking the presented sequence was displayed on the left and right to cue the respective hand that should execute the sequence. A red box appeared on the other side indicating that the hand should remain at rest on the keyboard. The instructional stimulus remained on the screen for 2.7s, after which five white asterisks were presented as a ‘go’ signal. Each instructed sequence was executed either four times in a row (in the pre-test and post-test) or three times (during training and fMRI acquisition), and each execution was individually triggered with a ‘go’ signal. There was a 500ms interval between consecutive sequence executions. For the remainder of the manuscript, we define a single ‘trial’ as the set of three or four consecutive executions of the same sequence.

The objective of the sequence task was to perform the five-finger presses as quickly as possible with minimal errors. Each finger was required to exceed a force of 2.5 N, while all other digits applied forces below 2.2 N. After each finger press, the corresponding asterisk changed color to provide feedback about whether the individual press was: ‘correct’ (green), ‘incorrect’ (red), or ‘too hard’ (yellow; that is, greater than the upper limit set for the task, 8.9 N). Execution time (ET) was measured as the duration between onset of the first press and release of the last press, and error rate was defined as the percentage of sequences that contained one or more incorrect finger presses. Throughout behavioral training, a constant error rate was encouraged by instructing participants to speed up if error rate was lower than 20% and slow down if it was higher.

After each sequence execution, participants saw a brief feedback (0.8s): one green asterisk (= 1 point) indicated that all five presses of the sequence were executed correctly; three green asterisks (= 3 points) meant that the sequence was executed correctly and with ≥20% faster ET than the average of the previous run; one blue asterisk specified that the sequence was performed 20% slower than the average of the previous run (= 0 points); and one red asterisk signified that one or more errors were made (= −1 point). Participants received a financial bonus according to their point score.

### Experimental Design

All participants completed 4 study phases (**Fig. 1**): a pre-test (day 1) in which baseline performance for twelve 5-finger sequences was evaluated for both hands; a 4-day training phase (days 2-5) during which participants repeatedly practiced the same four sequences with either the left or right hand for approximately an hour (which was coupled with 25 min of tDCS from the onset of training); a post-test (day 6), which was conducted in the same manner as the pre-test; and an MRI session (day 7).

The pre-test started with a short practice run with 4 trials or 16 executions (4 executions of two easy sequences per hand) to familiarize participants with the task. During the pre-test, participants performed all 12 sequences (4 to-be-trained and 8 untrained) with both left and right hands. Each hand was required to perform 2 trials per sequence with 4 executions per trial (i.e. 8 total executions). The pre-test consisted of eight runs with 24 trials per hand, which resulted in a total of 96 executions per hand. Within the first four runs, the order of sequences and hands was randomly permuted, and the order was reversed in the second half to counterbalance possible learning effects.

We assigned subjects pseudo-randomly to one of four stimulation groups (sham, unihemispheric, bihemispheric, or reversed-polarity bihemispheric tDCS groups) and to training cohort (left or right hand training), such that group differences in pre-test performance were minimized (33). ANOVA across tDCS groups revealed no difference in baseline task ET (*F*_(3,60)_ = 0.158, *p* = 0.923) (see **Table S1** for individual group means), and there was also no significant pair-wise difference between any two stimulation groups (all *t* < 1.10, *p* = 0.273).

To ensure that both participants and experimenter were blind to tDCS assignment, the randomisation was performed by a JD, who was not involved in data collection. The experimenter (SW) only knew the hand training cohort and the electrode arrangement, but not whether the participant received real or sham stimulation, which was determined by a randomized code entered into the tDCS machine at the beginning of each session. Accordingly, 66.7% of the sham participants (14/21) had bihemispheric electrode arrangement and 33.3% had unihemispheric electrode arrangement. No significant behavioral differences were found between these two subsets of the sham group on post-test performance (*F*_(1,16)_<1.454, *p*>0.245; ANCOVA using pre-test as covariate) for trained/untrained hand and trained/untrained sequence ET.

During the four training days, participants practiced 4 of the 12 sequences with either their left or right hand. A session consisted of 16 runs with 2 trials per sequence each.

Therefore, participants performed 128 trials (384 sequence executions) per day. On the day following the final session of tDCS-coupled training, a post-test that had exactly the same structure as the pre-test was administered.

### Transcranial direct current stimulation

tDCS was administered via a bihemispheric or unihemispheric montage. In the unihemispheric montage, we placed the anode over contralateral M1 and cathode over ipsilateral supraorbital ridge. For the bihemispheric montage, we positioned the anode over contralateral and the cathode above ipsilateral M1, with the reversed-polarity montage involving the opposite polarity. The hand area of M1 was localized as the position where single-pulse suprathreshold TMS (Magstim BiStim2 with a 5-cm figure-of-eight coil positioned tangentially to the skull at a 45° angle) evoked a visible twitch in the contralateral first dorsal interosseus muscle. tDCS was administered over four consecutive training days for the first 25 min of an approximately 60-min session of sequence training. A current of 2 mA was delivered using a neuroConn DC-stimulator PLUS (http://www.neuroconn.de/dc-stimulator_plus_en/) through saline-soaked 35 cm^2^ electrodes.

### Behavioral analysis

There was no significant difference between the tDCS groups in terms of error rate during the pre-test (F_(3,60)_ = 0.252, p = 0.860; averaged across training group and hand) or training (F_(3,60)_ = 1.082, p = 0.364; averaged across day). We only found a significant effect of tDCS on error rate in the post-test, with tDCS recipients tending to be more accurate than sham (F_(3,60)_ = 3.086, p = 0.0339; averaged across training group and hand).

To adjust ET for the different error rates, we calculated the median ET for each run, sequence, and hand over all (correct and incorrect) trials. For this calculation, ET for the incorrect trial was replaced with the maximum ET of that group of trials, thereby penalizing inaccurate performance.

For statistical comparison between pairs of groups, we used an ANCOVA in which the error-adjusted ET at pre-test was used as a covariate to account for prior inter-individual differences in performance. This procedure effectively subtracts from each training/post-test measurement the best prediction based on the pre-test measurement. Compared to simply subtracting the pre-test measurement, this method is less susceptible to noise induced by the larger variability in the pre-test measurements. The ANCOVA included ‘tDCS group’ and ‘hand training cohort’ as between-subject factors. The threshold for all statistical comparisons was *p* < 0.05. All data presented in the text and figures are represented as mean ± standard error of the mean (SEM).

### fMRI data acquisition

All 42 bihemispheric tDCS recipients (sham and real) underwent fMRI scanning one day after the post-test. At this point, at least 48h had elapsed since the final tDCS application. Therefore any group differences should be due to long-term neuroplastic changes induced by tDCS; immediate online effects of tDCS (34-36) should have been washed out.

The fMRI session consisted of 8 runs comprised of 24 randomly ordered trials (3 per trial type—4 sequences x 2 hands—with 3 sequence executions per trial, yielding 72 total executions per run). Each trial consisted of a cueing phase (2.7 s, 1 TR), followed by 3 executions of the cued sequence, triggered 3.6s apart. A trial, therefore, lasted 13.5s (5 TRs). Each sequence execution had to be completed within 2.8s to allow for a 0.8s feedback phase.

Baseline BOLD activation was measured during 8 randomly interspersed rest phases of 13.5s. To monitor for mirror activity on the non-moving hand, participants were required to keep all ten digits on the keyboard and to generate a small baseline force of ~0.5N at all times. Functional images were acquired using a 3T Siemens Trio MRI machine, with a 32-channel head coil. A 2D echo-planar sequence with a TR of 2.72s was used to acquire the functional volumes (8 runs, 159 volumes per run, 32 interleaved slices with 2.7 mm thickness) in an interleaved manner (3mm gap, and 2.3×2.3 mm^2^ in-plane resolution). The volumes were acquired in an oblique orientation with a 45° tilt angle from the AC-PC line—this slice prescription provided coverage of motor regions on the dorsal surface of the cortex, as well as the superior cerebellum and basal ganglia, but excluded inferior prefrontal and inferior and anterior temporal lobes. For full details of the fMRI acquisition, see (32).

### Behavior during scanning

To match behavioral performance during scanning, participants were instructed to produce the sequence within 1.3s and as accurately as possible. This ET was the fastest that subjects across all groups could achieve with both trained and untrained hands with high accuracy (~90% correct). Additionally, throughout training, force levels of the sequences were kept similar by imposing a maximal force threshold. No force level feedback was provided during fMRI scanning. Despite relatively well-matched performance, there were small significant differences between groups (see ‘Confound analysis’).

### fMRI analysis

The analysis of the fMRI data is described in detail inWiestler, Waters-Metenier and Diedrichsen (32), which reports the results from the sham group. After standard pre-processing (correction for slice acquisition, motion realignment, and coregistration to the individual anatomical image), we used a first-level linear model implemented in SPM8 (50) to estimate the activation for each of the four sequences for each hand. The design matrix consisted of a regressor for each hand and sequence type and an intercept for each run. The regressor was modelled as a boxcar function of 10.8s beginning with the first go cue of the trial, which was then convolved with a standard hemodynamic response function. From the estimated regression weights, we computed the percent signal change compared to rest for each hand, averaged across the four sequences.

For the group analysis, we reconstructed the cortical surface of each individual participant using Freesurfer (51), which permits the extraction of the white-grey matter surface and pial surface from anatomical images. The functional data was then projected onto each individual surface by averaging for each surface node the voxels that lay between the white-grey matter and pial surfaces.

The individual surfaces were then aligned to a shared spherical template for left and right hemispheres. This allowed us to flip the results for the right hand trained cohort, such that data from the trained hemisphere—the hemisphere contralateral to the trained hand—was displayed on the right group hemisphere, and the data from the untrained hemisphere on the left. We then conducted *t*-tests between sham and tDCS groups for each surface vertex, using an uncorrected height threshold of *T*_(1,39)_ = 2.71, *p* = 0.005. Family-wise error was controlled by calculating the critical size of the largest suprathreshold cluster that would be expected by chance, using Gaussian Field theory as implemented in the fmristat package (52). Results were displayed using the 3D-visualisation software Caret (53).

For the profile plots shown in **Fig.3a,b**, we defined a line running from the posterior parietal cortex through the hand area of M1 to the anterior tip of Brodman area 6 (premotor cortex). We then averaged activity over the area 1.5cm above and below that line (purple area in **Fig.3d**).

We also conducted a region of interest (ROI) analysis for the cortical motor regions. They included the hand region of primary motor cortex (M1, Brodman area 4); primary somatosensory cortex (S1, Brodman area 1-3); dorsal premotor cortex (PMd); supplementary motor areas (SMA/pre-SMA); and superior parietal lobe, divided into an anterior (intra-parietal sulcus, IPS) and posterior (occipito-parietal junction, OPJ) aspect. We defined all ROIs on the symmetric group template and subsequently projected this into individual data space via the respective individual surface.

The average activity in these ROIs was submitted to a ANOVA to calculate differences between tDCS groups (factor tDCS) and interactions with training hand (factor ‘hand’) and hemisphere (factor ‘hemisphere’). Because there were no significant functional differences between left and right hand training cohorts (F_(1,36)_ < 2.21, p > 0.15, for all 6 ROIs, even without correction), or interactions with any other factor, we analysed the functional data averaged over left and right hand training cohorts.

### Confound analysis

Despite our efforts to match behavior during fMRI acquisition across the tDCS groups, there were some significant differences between the groups (**Table 2**). To account for these parameters in our statistical model, we performed ANCOVA analysis using each behavioral parameter as a covariate for every ROI with a significant percent signal change difference due to tDCS (i.e. M1 and S1).

Another potential confound that could lead to increased ipsilateral activation is mirroring. Mirroring refers to the phenomenon whereby muscles of the non-moving hand are activated simultaneously with those of the moving hand (54). Such movements are typically visible in pathological states (e.g. stroke), but are also present and measurable in healthy populations. During fMRI, participants were required to rest the passive hand on the keyboard while the active hand was executing sequences. Mirroring was parameterized using the range of forces on the passive hand across the time course of a trial, the associated standard deviation, and the correlation between the force traces for matching digits of the passive and active hand. There was no significant difference between tDCS groups for any of these measures (**Table S2**).

## Acknowledgements

The research was supported by grants from the Wellcome trust (094874/Z/10/Z) and James McDonnell foundation, both to JD, and a PhD studentship from the Brain Research Trust (6CHB) to SW. The Wellcome Trust Centre for Neuroimaging is supported by core funding from the Wellcome Trust 091593/Z/10/Z).

## Supplemental Information

### Quantification of subject blindedness and perceptual side effects of tDCS

Subsequent to each session of tDCS administration, participants underwent a previously designed battery to characterize tDCS effects (33), with the addition of questions about perceptual side effects. Specifically, participants were asked to rate the experience of tingling, pain, burning, itching, dizziness, and mental fatigue (1:10) and then to describe (in min) the duration of the effect. As can be seen in **Fig. S2**, tDCS recipients tended to exhibit slightly higher intensities of perceptual properties of tDCS—especially in terms of tingling, burning and itching—however, none of these variables showed a significant between-group difference (all *t*_(62)_ < 1.446, *p* > 0.153), or exceeded level 4 (out of a maximal rating of 10) for any of the 64 subjects.

However, the groups tended to differ on how long they experienced these side effects. Averaged across the three tDCS groups (unihemispheric, bihemispheric and reversed-polarity bihemispheric), subjects reported significantly longer durations of tingling (*t*_(62)_ = 2.553, *p* = 0.013), burning (*t*_(62)_ = 2.492, *p* = 0.015), and itching (*t*_(62)_ = 2.833, *p* = 0.006), and a tendency for longer duration of pain (*t*_(62)_ =1.806, *p* = 0.076). There were no differences in the duration of dizziness (*t*_(62)_ = 1.051, *p* = 0.298) or mental fatigue (*t*_(62)_ = 0.671, *p* = 0.505) between sham and tDCS groups. However, as observed previously (33), participants experienced no significant differences in overall discomfort, perceived tDCS intensity, or distraction due to tDCS (**Table 2**). Moreover, χ^2^ - goodness of fit tests showed that there were no differences in the detectability of tDCS assignment between tDCS groups and sham (**Table 2**). These findings collectively suggest that, despite slight perceptual differences, tDCS blinding methods were sufficient. This is congruent with other recent evidence (55), including a study that investigated over twice as many subjects at 2 mA (56).

### Assessment of behavioral side effects of tDCS

In addition to the main training task (the sequence task described above), we tested all bihemispheric tDCS and sham recipients on two additional tasks of manual skill during the pre-test and post-test: individuation (the ability to move the digits separately) and configuration execution (the skill of pressing certain digits at the same time while keeping coactivation of unintended digits minimal)—for full details of these tasks, see Waters-Metenier, Husain, Wiestler and Diedrichsen (33). Neither bihemispheric tDCS group exhibited any side effects from tDCS-coupled sequence training on either of these skills (**Fig. S2**), and there were no significant differences between the two bihemispheric groups and sham (all *F*_(2,35)_<2.092, p>0.139; using ANCOVA correction with pre-test performance). Therefore, as in our previous work, we did not find any adverse trade-offs between using tDCS to facilitate manual motor skill learning and performance on untrained tasks (33).

**Supplemental Figure S1:**
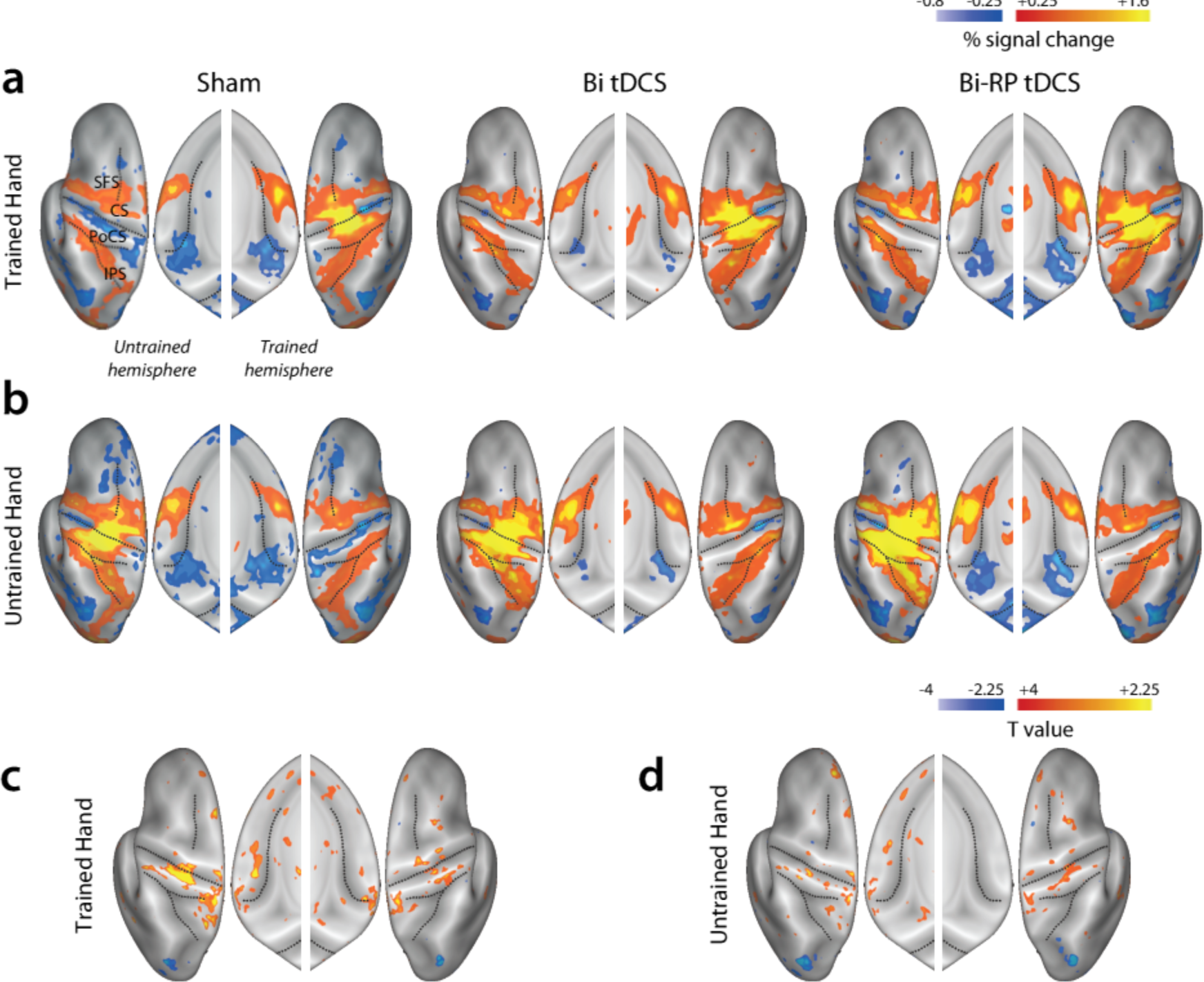
Bihemispheric tDCS groups exhibit more activity in bilateral sensorimotor areas relative to sham. **(a)** Activation maps for the trained and untrained hemispheres for the sham, bihemispheric (Bi tDCS), and reversed-polarity (Bi-RP tDCS) groups averaged across hand training cohort. While the sham group exhibited ipsilateral deactivation (typically found in fMRI studies of unilateral movements), this pattern was not found for either bihemispheric tDCS group. Additionally, tDCS groups exhibited higher contralateral activation. **(b)** Corresponding maps for sequences executed with the untrained hand. **(c,d)** Difference T-maps for the activation in the tDCS groups relative to sham for the trained (**c**) and untrained (**d**) hands. Abbreviations: CS = central sulcus; IPS = intraparietal sulcus; PoSC = postcentral sulcus; SFS = superior frontal sulcus.

**Supplemental Figure S2:**
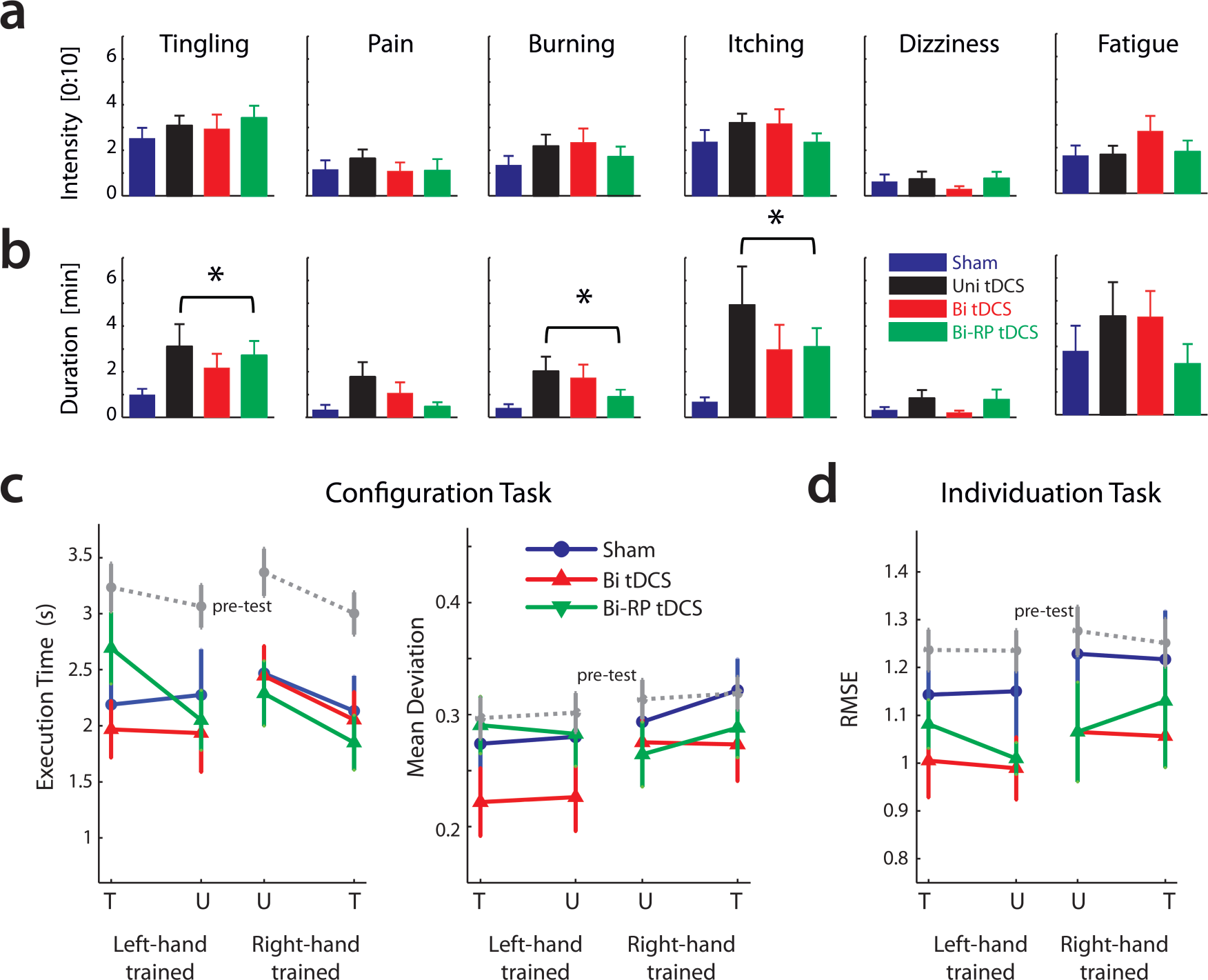
Online perceptual side effects of tDCS. We monitored the potential online side effects of tDCS in order to evaluate perceptual differences between tDCS groups and sham. **(a)** For the intensity level of all six effect types, there were no significant differences across tDCS groups. **(b)** However, the differences between tDCS and sham tended to be greater for the duration of these side effects and significant differences were observed for burning and itching *(indicated by asterisks).* Note that differences in duration of tingling and pain percepts were also nearly significantly different. **(c)** Post-test performance of the configuration task speed *(left panel)* and mean deviation *(right panel)* for the sham *(blue),* bihemispheric *(red),* and reversed-polarity bihemispheric *(green)* groups. **(d)** Post-test performance of individuation RMSE. tDCS recipients experienced no adverse behavioral effects as a result of tDCS-coupled sequence training.

**Supplemental Table S1:** Participant demographic and psychological variables. To evaluate differences in sex and detectability of tDCS status, all three tDCS groups were individually compared to sham using a^χ^2^^ - goodness of fit test. For all other parameters, mixed-effects AN OVA was calculated with the between factor tDCS. All variables were averaged across left and right hand training cohorts.

| Group | Sham | Uni tDCS | Bi tDCS | Bi-RP tDCS | Over all 4 groups |  |
| --- | --- | --- | --- | --- | --- | --- |
| N (LH-trained; RH-trained) | 21 (11;10) | 15 (8; 7) | 14 (7; 7) | 14 (7; 7) |  |  |
| Sex (Female : Male) | 13 : 8 | 7 : 8 | 6 : 8 | 9 : 5 | $\chi^2_{(1)}$ | $p$ |
|  |  |  |  |  | 0.66 | 0.42 |
| **Detectability of tDCS status** |  |  |  |  |  |  |
| tDCS status: (Yes: No) | 15 : 6 | 11 : 4 | 10 : 4 | 11 : 3 | 15:6 | 32:11 |
| | | | | | $\chi^2_{(1)}$ | $p$ |
|  |  |  |  |  | 0.06 | 0.80 |
| **tDCS perception** | | | | | $F_{(3,60)}$ | $p$ |
| **Averaged across all training days** |  |  |  |  |  |  |
| Discomfort (0:10) | 2.32 ± 0.46 | 2.72 ± 0.39 | 3.30 ± 0.54 | 2.88 ± 0.37 | 0.846 | 0.474 |
| Perceived Intensity (0:10) | 2.81 ± 0.40 | 3.28 ± 0.44 | 2.84 ± 0.62 | 3.91 ± 0.41 | 1.164 | 0.331 |
| Distraction due to tDCS (0:10) | 1.07 ± 0.31 | 1.80 ± 0.38 | 1.63 ± 0.43 | 1.57 ± 0.39 | 0.825 | 0.485 |
| **Demographics** | | | | | $F_{(3,60)}$ | $p$ |
| Age | 22.19 ± 0.95 | 24.00 ± 1.59 | 22.93 ± 0.99 | 22.43 ± 0.82 | 0.518 | 0.671 |
| Handedness (Edinburgh) | 88.09 ± 1.87 | 84.33 ± 4.57 | 77.5 ± 4.15 | 82.14 ± 3.95 | 1.661 | 0.185 |
| Previous motor training (hrs) | 640 ± 293 | 872 ± 396 | 447 ± 246 | 486 ± 306 | 0.336 | 0.800 |
| **Baseline motor performance** | | | | | $F_{(3,60)}$ | $p$ |
| Baseline ET<sub>A</sub> | 2.36 ± 0.065 | 2.33 ± 0.15 | 2.45 ± 0.10 | 2.32 ± 0.069 | 0.159 | 0.924 |
| **Psychological measures** | | | | | $F_{(3,60)}$ | $p$ |
| **Averaged across all days** |  |  |  |  |  |  |
| Sleep Hours | 7.29 ± 0.21 | 6.94 ± 0.22 | 7.49 ± 0.19 | 7.49 ± 0.21 | 1.393 | 0.254 |
| Sleep Quality (0:10) | 7.28 ± 0.26 | 6.97 ± 0.19 | 7.89 ± 0.24 | 7.27 ± 0.26 | 2.155 | 0.102 |
| Alertness (0:10) | 6.77 ± 0.35 | 6.62 ± 0.51 | 7.21 ± 0.55 | 7.12 ± 0.24 | 0.413 | 0.744 |
| Attention (0:10) | 7.65 ± 0.28 | 7.56 ± 0.34 | 8.18 ± 0.32 | 7.04 ± 0.52 | 1.490 | 0.226 |
| Task Difficulty (0:10) | 4.56 ± 0.45 | 5.43 ± 0.42 | 4.71 ± 0.37 | 5.16 ± 0.30 | 0.993 | 0.403 |
| **Averaged across final training day and post-test** |  |  |  |  |  |  |
| Semantic Recall (% correct) | 73.21 ± 4.59 | 88.33 ± 3.33 | 96.43 ± 1.57 | 93.75 ± 2.54 | 7.362 | 0.0003 |

**Supplemental Table S2:** Mirroring of finger forces during fMRI. Table shows mean (±standard error) of the average force range (maximum − minimum) and standard deviation of the passive (non-moving) hand. Results were split depending on whether the trained hand or the untrained hand was the passive hand. The correlation between the forces was calculated between the force time series of the matching digits of the active and passive hands. The last column indicates the F-test comparing the 3 groups.

|  | Sham |  | Bi tDCS |  | RP-Bi tDCS |  | ANOVA across groups |  |
| --- | --- | --- | --- | --- | --- | --- | --- | --- |
| | mean | SE | mean | SE | mean | SE | $F_{(2,39)}$ | $p$ |
| **<u>Trained Hand</u>** |  |  |  |  |  |  |  |  |
| Force range [N] | 0.13 | (0.01) | 0.14 | (0.01) | 0.15 | (0.03) | 0.356 | 0.703 |
| Standard deviation [N] | 0.03 | (0.00) | 0.03 | (0.00) | 0.04 | (0.01) | 0.497 | 0.612 |
| Correlation of force traces | 0.07 | (0.002) | 0.07 | (0.05) | 0.03 | (0.03) | 0.387 | 0.682 |
| **<u>Untrained Hand</u>** |  |  |  |  |  |  |  |  |
| Force range [N] | 0.12 | (0.01) | 0.12 | (0.01) | 0.13 | (0.02) | 0.293 | 0.745 |
| Standard deviation [N] | 0.03 | (0.00) | 0.03 | (0.00) | 0.03 | (0.01) | 0.260 | 0.773 |
| Correlation of force traces | 0.09 | (0.02) | 0.05 | (0.03) | 0.02 | (0.03) | 1.981 | 0.154 |

## References

1. Di Pino G, et al. (2014) Modulation of brain plasticity in stroke: a novel model for neurorehabilitation. Nature reviews. Neurology 10(10):597–608.

2. Perez MA & Cohen LG (2009) Interhemispheric inhibition between primary motor cortices: what have we learned? The Journal of physiology 587(Pt 4):725–726.

3. Chen R, Cohen LG, & Hallett M (1997) Role of the ipsilateral motor cortex in voluntary movement. The Canadian journal of neurological sciences. Le journal canadien des sciences neurologiques 24(4):284–291.

4. Murase N, Duque J, Mazzocchio R, & Cohen LG (2004) Influence of interhemispheric interactions on motor function in chronic stroke. Annals of neurology 55(3):400–409.

5. Takeuchi N, Oouchida Y, & Izumi S (2012) Motor control and neural plasticity through interhemispheric interactions. Neural plasticity 2012:823285.

6. Hummel FC & Cohen LG (2006) Non-invasive brain stimulation: a new strategy to improve neurorehabilitation after stroke? Lancet Neurol 5(8):708–712.

7. Nitsche MA (2011) Beyond the target area: remote effects of non-invasive brain stimulation in humans. The Journal of physiology 589(Pt 13):3053–3054.

8. Nitsche MA & Paulus W (2000) Excitability changes induced in the human motor cortex by weak transcranial direct current stimulation. The Journal of physiology 527 Pt 3:633–639.

9. Nitsche MA, et al. (2003) Level of action of cathodal DC polarisation induced inhibition of the human motor cortex. Clin Neurophysiol 114(4):600–604.

10. Vines BW, Cerruti C, & Schlaug G (2008) Dual-hemisphere tDCS facilitates greater improvements for healthy subjects' non-dominant hand compared to uni-hemisphere stimulation. BMC Neurosci 9:103.

11. Krause B, Marquez-Ruiz J, & Kadosh RC (2013) The effect of transcranial direct current stimulation: a role for cortical excitation/inhibition balance? Frontiers in human neuroscience 7:602.

12. Bolognini N, et al. (2011) Neurophysiological and behavioral effects of tDCS combined with constraint-induced movement therapy in poststroke patients. Neurorehabil Neural Repair 25(9):819–829.

13. Lefebvre S, et al. (2014) Single Session of Dual-tDCS Transiently Improves Precision Grip and Dexterity of the Paretic Hand After Stroke. Neurorehabil Neural Repair 28(2):100–110.

14. Nitsche MA & Paulus W (2011) Transcranial direct current stimulation--update 2011. Restorative neurology and neuroscience 29(6):463–492.

15. Williams JA, Pascual-Leone A, & Fregni F (2010) Interhemispheric modulation induced by cortical stimulation and motor training. Phys Ther 90(3):398–410.

16. Takeuchi N & Izumi S (2012) Maladaptive plasticity for motor recovery after stroke: mechanisms and approaches. Neural plasticity 2012:359728.

17. Zimerman M, et al. (2012) Modulation of training by single-session transcranial direct current stimulation to the intact motor cortex enhances motor skill acquisition of the paretic hand. Stroke 43(8):2185–2191.

18. Batsikadze G, Moliadze V, Paulus W, Kuo MF, & Nitsche MA (2013) Partially non-linear stimulation intensity-dependent effects of direct current stimulation on motor cortex excitability in humans. The Journal of physiology 591(Pt 7):1987–2000.

19. O'Shea J, et al. (2014) Predicting behavioural response to TDCS in chronic motor stroke. NeuroImage 85 Pt 3:924–933.

20. Rahman A, et al. (2013) Cellular Effects of Acute Direct Current Stimulation: Somatic and Synaptic Terminal Effects. The Journal of physiology.

21. Allison JD, Meador KJ, Loring DW, Figueroa RE, & Wright JC (2000) Functional MRI cerebral activation and deactivation during finger movement. Neurology 54(1):135–142.

22. Wiestler T & Diedrichsen J (2013) Skill learning strengthens cortical representations of motor sequences. eLife 2:e00801.

23. Diedrichsen J, Wiestler T, & Krakauer JW (2013) Two distinct ipsilateral cortical representations for individuated finger movements. Cerebral cortex (New York, N.Y.: 1991) 23(6):1362–1377.

24. Ganguly K, et al. (2009) Cortical representation of ipsilateral arm movements in monkey and man. The Journal of neuroscience: the official journal of the Society for Neuroscience 29(41):12948–12956.

25. Verstynen T, Diedrichsen J, Albert N, Aparicio P, & Ivry RB (2005) Ipsilateral motor cortex activity during unimanual hand movements relates to task complexity. J Neurophysiol 93(3):1209–1222.

26. Reis J & Fritsch B (2011) Modulation of motor performance and motor learning by transcranial direct current stimulation. Curr Opin Neurol 24(6):590–596.

27. Mahmoudi H, et al. (2011) Transcranial direct current stimulation: electrode montage in stroke. Disabil Rehabil 33(15–16):1383–1388.

28. Karok S & Witney AG (2013) Enhanced motor learning following task-concurrent dual transcranial direct current stimulation. PloS one 8(12):e85693.

29. Lindenberg R, Nachtigall L, Meinzer M, Sieg MM, & Floel A (2013) Differential Effects of Dual and Unihemispheric Motor Cortex Stimulation in Older Adults. The Journal of neuroscience: the official journal of the Society for Neuroscience 33(21):9176–9183.

30. Sehm B, Kipping J, Schafer A, Villringer A, & Ragert P (2013) A Comparison between Uni- and Bilateral tDCS Effects on Functional Connectivity of the Human Motor Cortex. Frontiers in human neuroscience 7:183.

31. Naros G, et al. (2016) Enhanced motor learning with bilateral transcranial direct current stimulation: Impact of polarity or current flow direction? Clin Neurophysiol 127(4):2119–2126.

32. Wiestler T, Waters-Metenier S, & Diedrichsen J (2014) Effector-independent motor sequence representations exist in extrinsic and intrinsic reference frames. The Journal of neuroscience: the official journal of the Society for Neuroscience 34:5054–5064.

33. Waters-Metenier S, Husain M, Wiestler T, & Diedrichsen J (2014) Bihemispheric Transcranial Direct Current Stimulation Enhances Effector-Independent Representations of Motor Synergy and Sequence Learning. Journal of Neuroscience 34(3).

34. Antal A, et al. (2014) Imaging artifacts induced by electrical stimulation during conventional fMRI of the brain. NeuroImage 85 Pt 3:1040–1047.

35. Mielke D, et al. (2013) Cathodal transcranial direct current stimulation induces regional, long-lasting reductions of cortical blood flow in rats. Neurological research.

36. Vernieri F, et al. (2010) Cortical neuromodulation modifies cerebral vasomotor reactivity. Stroke 41(9):2087–2090.

37. Johansen-Berg H, et al. (2002) The role of ipsilateral premotor cortex in hand movement after stroke. Proc Natl Acad Sci U S A 99(22):14518–14523.

38. Zimerman M, Heise KF, Gerloff C, Cohen LG, & Hummel FC (2014) Disrupting the ipsilateral motor cortex interferes with training of a complex motor task in older adults. Cerebral cortex (New York, N.Y.: 1991) 24(4):1030–1036.

39. Truong DQ, et al. (2014) Clinician Accessible Tools for GUI Computational Models of Transcranial Electrical Stimulation: BONSAI and SPHERES. Brain stimulation.

40. Kwon YH & Jang SH (2011) The enhanced cortical activation induced by transcranial direct current stimulation during hand movements. Neurosci Lett 492(2):105–108.

41. Stagg CJ, et al. (2012) Cortical activation changes underlying stimulation-induced behavioural gains in chronic stroke. Brain: a journal of neurology 135(Pt 1):276–284.

42. Baudewig J, Nitsche MA, Paulus W, & Frahm J (2001) Regional modulation of BOLD MRI responses to human sensorimotor activation by transcranial direct current stimulation. Magn Reson Med 45(2):196–201.

43. Kwon YH, et al. (2008) Primary motor cortex activation by transcranial direct current stimulation in the human brain. Neurosci Lett 435(1):56–59.

44. Jang SH, et al. (2009) The effect of transcranial direct current stimulation on the cortical activation by motor task in the human brain: an fMRI study. Neurosci Lett 460(2):117–120.

45. Kim CR, et al. (2012) Modulation of cortical activity after anodal transcranial direct current stimulation of the lower limb motor cortex: a functional MRI study. Brain stimulation 5(4):462–467.

46. Stagg CJ, et al. (2009) Modulation of movement-associated cortical activation by transcranial direct current stimulation. Eur J Neurosci 30(7):1412–1423.

47. Stagg CJ, et al. (2009) Polarity-sensitive modulation of cortical neurotransmitters by transcranial stimulation. The Journal of neuroscience: the official journal of the Society for Neuroscience 29(16):5202–5206.

48. Diedrichsen J & Kornysheva K (2015) Motor skill learning between selection and execution. Trends in cognitive sciences 19(4):227–233.

49. Oldfield RC (1971) The assessment and analysis of handedness: the Edinburgh inventory. Neuropsychologia 9(1):97–113.

50. Friston KJ, Ashburner J, Kiebel SJ, Nichols TE, & Penny WD eds (2007) Statistical Parametric Mapping: The Analysis of Functional Brain Images (Academic Press).

51. Dale AM, Fischl B, & Sereno MI (1999) Cortical surface-based analysis. I. Segmentation and surface reconstruction. NeuroImage 9(2):179–194.

52. Worsley KJ, et al. (1996) A unified statistical approach for determining significant signals in images of cerebral activation. Human brain mapping 4(1):58–73.

53. Van Essen DC, et al. (2001) Mapping visual cortex in monkeys and humans using surface-based atlases. Vision research 41(10–11):1359–1378.

54. Beaule V, Tremblay S, & Theoret H (2013) Interhemispheric control of unilateral movement. Neural plasticity 2012:627816.

55. Kessler SK, Turkeltaub PE, Benson JG, & Hamilton RH (2012) Differences in the experience of active and sham transcranial direct current stimulation. Brain stimulation 5(2):155–162.

56. Russo R, Wallace D, Fitzgerald PB, & Cooper NR (2013) Perception of Comfort During Active and Sham Transcranial Direct Current Stimulation: A Double Blind Study. Brain stimulation doi: 10.1016/j.brs.2013.05.009. [Epub ahead of print].

